# Learning a non-neutral conditioned stimulus: place preference in the crab *Neohelice granulata*

**DOI:** 10.1101/2020.12.18.423472

**Authors:** Martín Klappenbach, Candela Medina, Ramiro Freudenthal

**Affiliations:** Instituto de Fisiología, Biología Molecular y Neurociencias; Universidad de Buenos Aires; CONICET; Ciudad autónoma de Buenos Aires; Argentina; Laboratory of Neural Plasticity and Memory, Institute of Biosciences, Biotechnology and translational Biology (iB3), Department of Physiology Molecular and Cellular Biology, University of Buenos Aires/CONICET, Ciudad Autónoma de Buenos Aires, Argentina

**Keywords:** Long-term memory, Place preference, Neohelice granulata, classical conditioning, Non-neutral conditioned stimulus

## Abstract

In the wild, being able to recognize and remember specific locations related to food sources and the associated attributes of landmarks is a cognitive trait important for survival. In the present work we show that the crab Neohelice granulata can be trained to associate a specific environment with an appetitive reward in a conditioned place preference task. After a single training trial, when the crabs were presented with a food pellet in the target quadrant of the training arena, they were able to form a long-term memory related to the event. This memory was evident at least 24 h after training and was protein-synthesis dependent. Importantly, the target area of the arena proved to be a non-neutral environment, given that animals initially avoided the target quadrant. In the present work we introduce for the first time an associative one-trial memory paradigm including a conditioned stimulus with a clear valence performed in a crustacean.

## Introduction

Contextual memory is a very important cognitive function, as it allows the storage and retrieval of specific cued-based locations which are of relevance to animals (Mery, 2013; Shettleworth, 2010). The crab Neohelice granulata lives in the intertidal flats of rias in Atlantic shores, where its populations form dense patches, with up to 70 burrows per m2 (Angeletti and Cervellini, 2015). Therefore, this species’ orientation skills are key components for successful navigation in this rich environment (Fathala and Maldonado, 2011). Several studies performed in maze and place preference arenas indicate that decapods are able to acquire and make use of learned information to orient themselves in rich environments (Davies et al., 2019; Huber et al., 2018; Tierney and Lee, 2011). In particular, Neohelice granulata is a strong model to study neurobiology of memory (Tomsic and Romano, 2013, Feld et al., 2020). In the present work we introduce for the first time a place preference long-term memory paradigm in which Neohelice granulata crabs are able to form an association between a striped identifiable quadrant (StQ) of a circular arena and an appetitive reinforcement, which led to an increase in their relative presence in the target quadrant 24 hours post-training. Interestingly, we show that the StQ is a non-neutral stimulus and that the initial amount of time spent on the first day in the quadrant can be modulated by the animal’s hunger state. This novel paradigm allows animals to form a long-term memory that can be modulated by the amount of time the contingency between stimuli takes place and is dependent on protein synthesis.

## Materials and Methods

### Animals

Adult male intertidal crabs (*Neohelice granulata*, formerly *Chasmagnathus granulatus*) were collected from water <1 m deep in narrow coastal inlets (rias) of San Clemente del Tuyú, Buenos Aires Province, Argentina. Only animals measuring 2.7-3.0 cm across the carapace and weighing ~17 g were selected to perform experiments and transported to the laboratory. Crabs were housed in plastic tanks (35 × 48 × 27 cm) filled to a depth of 2 cm with diluted marine water (Red sea fish pharm) with a salinity of 1.0%-1.4% and a pH of 7.4-7.6. The water was changed and the tank sanitized every 2 d. The housing room was maintained on a 12-h light-dark cycle (lights on from 07:00-19:00 h) and temperature between 22°C-24°C. Animals were food-restricted for 6-10 days depending on the experiment. These are opportunistic animals that can go through several days without feeding in the wild (Sarapio et al., 2017). Experiments are performed only in males to avoid disrupting the natural population or crabs since females carry the fecundated eggs in the first stages of development and capturing them might affect the size of the population. Another factor considered is that a natural population has many sources of variability and selecting male animals restricts variables such as size range. The reported research was conducted in accordance with the local regulations for the care and use of laboratory animals. All experiments were done in accordance to local regulations to minimize animal suffering and the number of animals used.

### Training and testing procedures

The conditioning apparatus comprises a cylindrical white PVC container with 30 cm tall wall which delimits a 31.5 cm circumference arena and it is shown in Figure 1. One quarter of the container wall has a black and white striped vertical pattern (StQ), in contrast with the rest of the arena’s wall, which is plain white. The floor of the arena is also plain white, with the exception of the striped quadrant delimitation in the container floor. For each experiment, the arena’s floor was covered to a depth of 0.5 cm with salt water. The containers were placed on top of a 130 cm stand to avoid visual contact between the experimenter and crabs were individually placed in each arena. A structural scaffold was placed 150 cm above an array of 8 arenas, with two high-frequency fluorescent lamps and two 922 Logitech cameras (each one recording a subset of 4 arenas) attached. Experimental groups were homogeneously distributed in all conditioning arenas and each arena’s orientation was rotated between trials, ensuring a distribution of the animal groups that takes into account the possible inhomogeneities of the experimental room (Figure 1).

**Figure 1.**
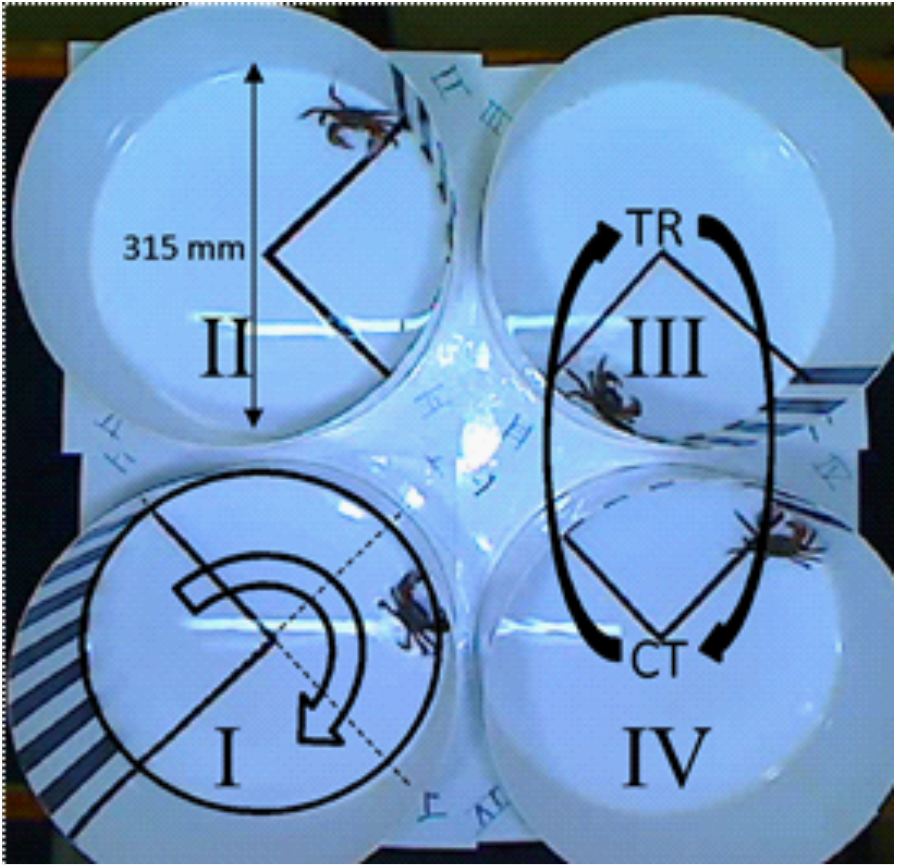
Arrange of a subset of four place preference arenas. Empty black arrow denotes the sense of rotation of the arena. Filled black arrows denoted the exchange of experimental groups between arenas among the experiment.

Video recordings of habituation (Hab), training (TR) and testing (TS) sessions were used to track each animal’s position within the arena manually and semi-automatically by Tracker 5.0.7, both in an offline manner (video analysis and modeling tool by Douglas Brown; https://physlets.org/tracker/). Heat maps of figures 2 and 4 where compiled using ANY-maze behavioural tracking software. The training session lasted up to 15 min and consisted of 5 min habituation to the arena, followed by a variable period of time during which the animal was presented with one pellet of food (Bottom fish, Labcon) at the bottom of the container in the center of the StQ. Then, animals were removed from the experimental arenas and individually housed in plastic containers, covered to a depth of 0.5 cm with seawater and kept inside dimly lighted drawers until the testing session, which took place 24 hours later. The testing session consisted of 5 min exposure to the conditioning arena without the presence of reward. The manual tracking consisted in evaluating the presence of the crab in the striped quadrant every 10 s for the 300 s duration of habituation or testing sessions. This variable was expressed as a percent of the 30 measurements in which the animal was present in StQ.

**Figure 2.**
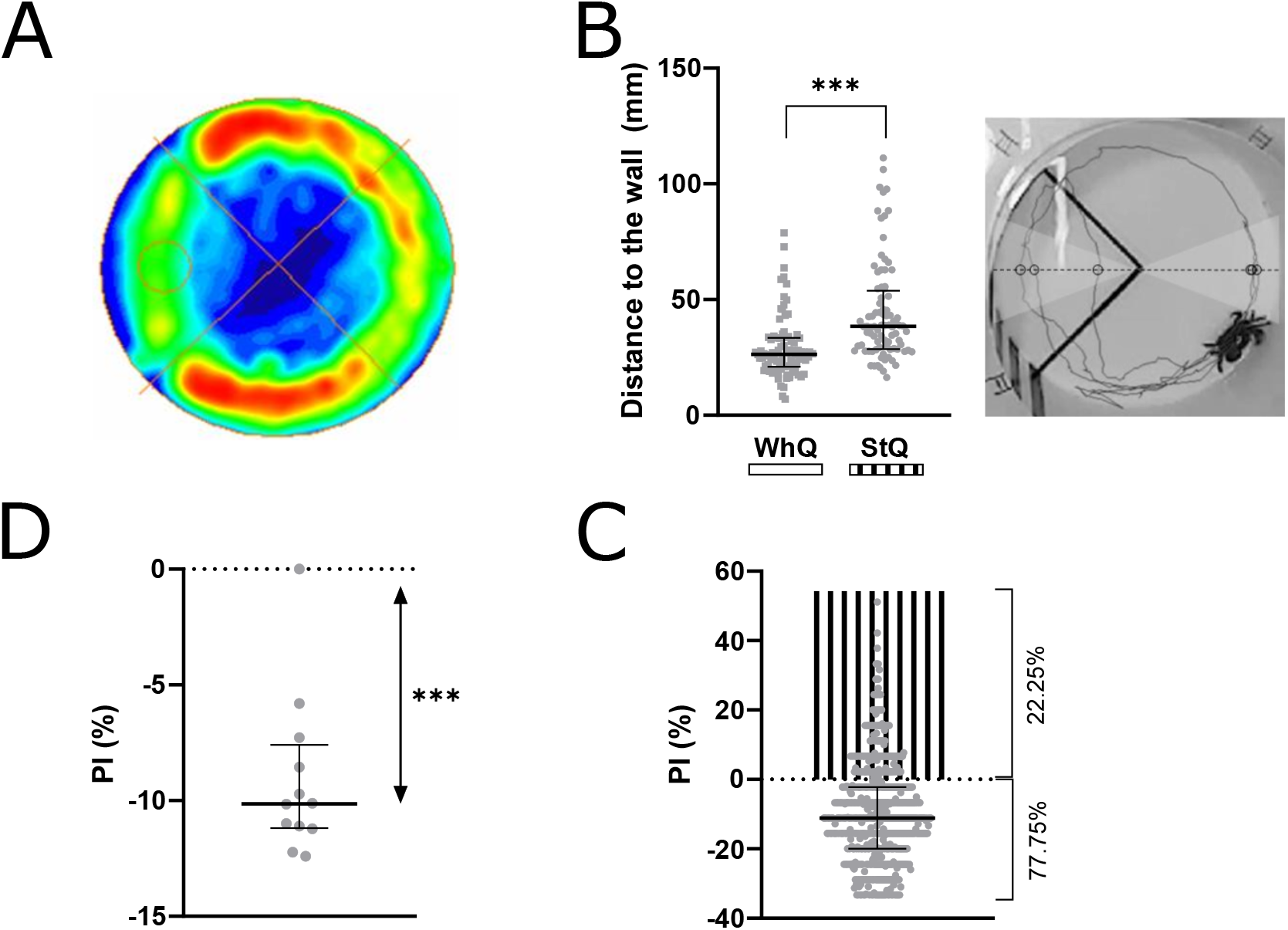
Crabs avoid the striped quadrant. (A) Heat map representing exploration distribution of animals during habituation session. The StQ is identified with a circle (n = 20). (B) Left: Distance to the wall in the StQ vs WhQ. Median ± interquartile ranges. Wilcoxon test between quadrants: W = −2788, p < 0.001. Right: circles represent intersection of a single crab trajectory with the bisector of each quadrant angle, denoting distance from the wall for both quadrants in the same axis. (C) StQ preference index (PI) 77.75% of animals spend more time in the StQ than the rest of the arena, while 22.25% of crabs present the opposite behavior, n=701. Median ± interquartile ranges. (D) PI calculated for each independent experiment (n = 12 experiments, 40-80 crabs/experiment; One sample Wilcoxon Signed Rank Test: W = −78, p < 0.001.

### Data analysis

Animals which displayed more than two standard deviations from the mean on habituation session were excluded from the analysis. This criterion was used for all experiments, except in the CS-training Figure, where all animals that showed times in the StQ higher than 25% were excluded. The StQ preference index (PI) was calculated as: PI = percentage of StQ time - (percentage of white quadrants time/3). Statistical analyses were performed using Graphpad prism software. Data were tested for normality using the Shapiro-Wilk test and analyzed with a Repeated Measures ANOVA (Holm-Sidak as posthoc test, when contrasts were pertinent), unless otherwise stated. Data was presented as mean ± standard errors mean (SEM), unless otherwise stated in the cases when it deviated from normality. One sample Wilcoxon Signed Rank Test was applied to compare the median of the sample with a hypothetical median (in this case, zero). When stated in Figures: * stands for p< 0.05, ** p< 0.01, *** p< 0.001.

### Drugs and injection procedure

The protein synthesis inhibitor cycloheximide (CHX, Sigma-Aldrich C7698) (Pedreira et al., 1995) was diluted in crustacean saline solution (450 mM NaCl, 15 mM CaCl2, 21 mM MgCl2, 10 mM KCl) (Hoeger and Florey, 1989) and administered systemically at a final dose of 40 μg/animal. Fifty microliters of vehicle (VHC) or drug solution were injected through the right side of the dorsal cephalothoraxic-abdominal membrane by means of a syringe needle fitted with a sleeve to control the depth of penetration to 4 mm, thus ensuring that the injected solution was released in the pericardial sac. Given that crabs lack an endothelial blood-brain barrier (Abbott, 1970),and that blood is distributed by a capillary system in the central nervous system (Sandeman, 1967), systemic injected drugs readily reach the brain.

## Results

### Animals explore the arena asymmetrically

During their first exposure to the arena (habituation session; Hab), crabs showed an inhomogeneous distribution of the exploratory behavior. In all four quadrants of the arena, animals spend most of the exploration time (5 min) in the close proximity of the walls (Figure 2A, B), suggesting thigmotaxis as a driver in their behavior. Nonetheless, the distance to the wall kept by the animals in the striped quadrant (StQ) is significantly larger than the distance shown in the exact opposite white quadrant (WhQ), indicating that the striped pattern is a relevant stimulus for these crabs (Figure 2B). The StQ preference index (PI) denotes the exploration time distribution of the animals in the arena. During habituation, 77.75% of the animals present a PI < 0, indicating they spend less time in the StQ than in the rest of the arena, while 22.25% present the opposite behavior (Figure 2C).

When the StQ PI during habituation was calculated for 12 independent experiments the resulting median is significantly lower than the null median expected for no preference (median = −10.14; p < 0.001, Figure 2D). This result indicates that the avoidance towards the StQ by the majority of the animals is the most frequent response to the initial presentation of the arena.

Considering the ever-changing relationship between an animal and its environment, we hypothesized that this initial avoidance towards the StQ could be affected by a number of factors, including the animal’s fasting period previous to the habituation session. An animal that has been food deprived will show increased drive for food seeking behaviors, and may show increased exploratory behaviors, with the potential to overcome mildly aversive stimuli as the stripes (Dethier, 1976). Fasting may also result in more robust learning experiences in crabs and other species (Klappenbach et al., 2017; Krashes et al., 2009). Effectively, we found a positive significant correlation between the number of days animals were food-deprived before their first encounter with the arena (i.e., habituation session) and the time spent by each animal in StQ (Figure 3A; r2 = 0.55; p < 0.01). We also found a positive significant correlation between the number of days the animals were food deprived and the fraction of animals preferring the StQ (Figure 3B; r2 = 0.49; p < 0.05). Lastly, this significant positive correlation was also evidenced in the quadrant PI (Figure 3C; r2 = 0.54; p < 0.01). These results point to an important modulating role of the animal’s motivational state by hunger in the exploration strategies in a novel environment.

**Figure 3.**
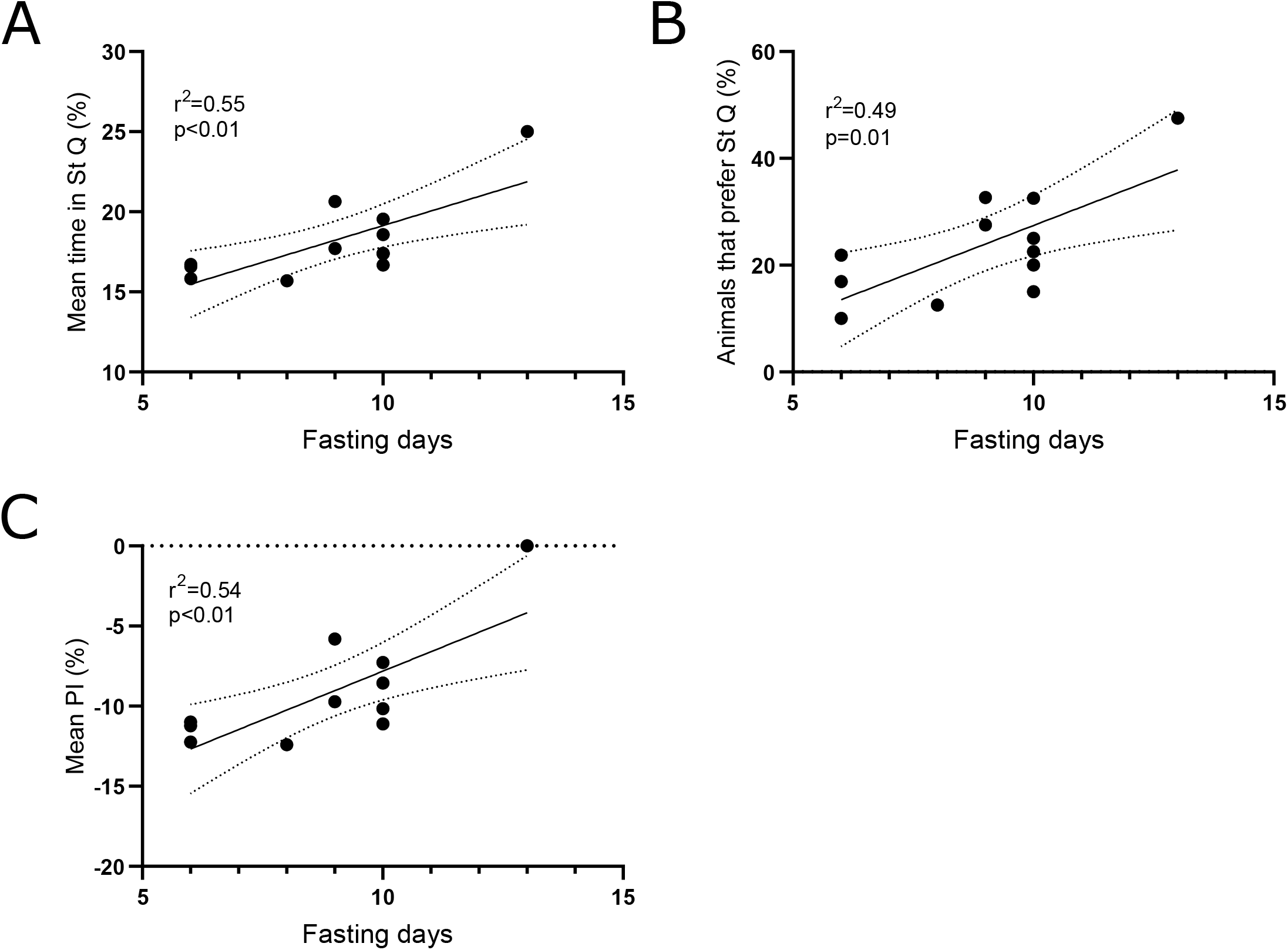
The time in the striped quadrant is modulated by the number of previous fasting days. (A) Mean percentage of time spent exploring StQ in animals fasting for 6 to 11 days. n = 12 independent experiments. (B) Percentage of animals which spend more time in the StQ when fasting for 6 to 11 days. n = 12 independent experiments. (C) Mean StQ PI in animals fasting for 6 to 11 days. n = 12 independent experiments.

### Seeking and feeding behavior in Neohelice granulata

Once crabs had gone through the habituation period in the arena, we randomly divided them into two groups. Animals from the trained group (TR) received one food pellet in the center of the StQ, to act as an appetitive reinforcement up until animals were removed, 10 min later. The other half of the animals remained in their arenas the same amount of time as the trained group but without the appetitive reinforcement, serving as control (CT). On average, animals from the TR group took 88.68 sec to find the pellet (Figure 4A) and this resulted in an effective reinforcement period of 497.3 sec (time spent in StQ eating, after finding the pellet; Figure 4B). Animals that experienced less than half of the mean effective reinforcement time were excluded from further analysis (identified with an “x” in Figures 4A and B). This could be due to presenting a longer period of time to find the food pellet, to abandoning it in order to further explore the environment or taking it with them away from the StQ. Nevertheless, this type of behavior was not usual, since after finding the pellet crabs spend 97.5% +/- 1.66 SEM of the time eating within StQ and 98.2% +/-1.7 SEM of this time they remain in the same position, as seen in Figure 4C. These characteristics of the crabs’ feeding behavior allowed a prolonged unforced association between the StQ and the appetitive reinforcement.

**Figure 4.**
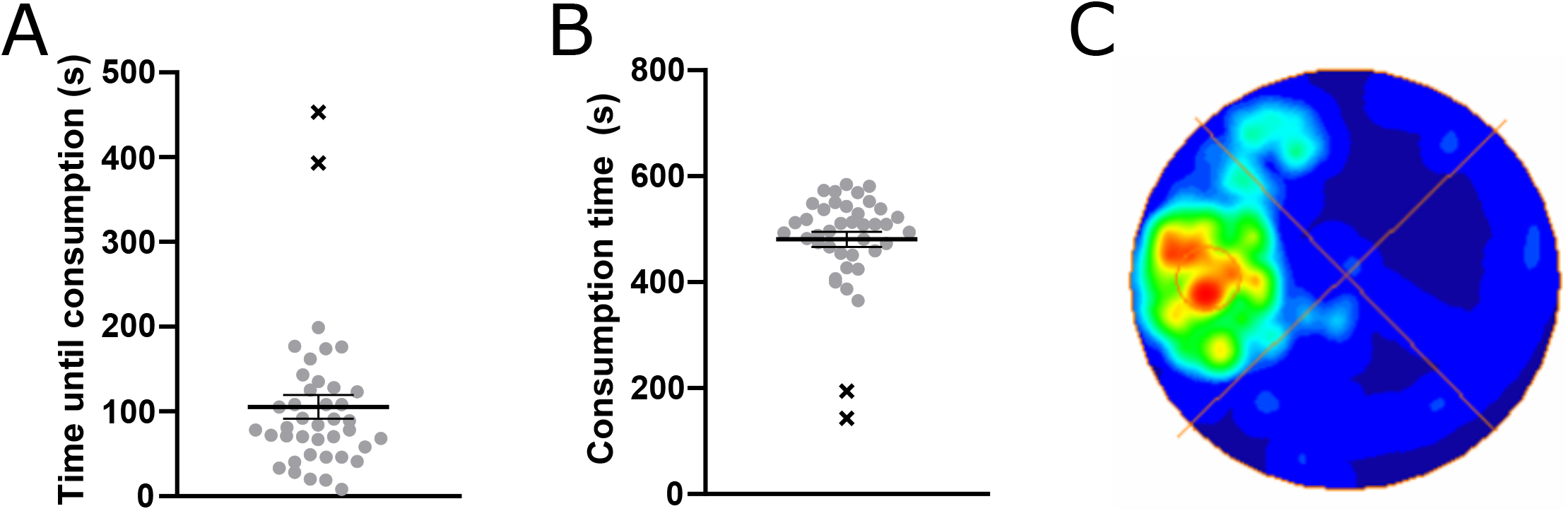
Seeking and feeding behavior in the arena. (A) Amount of time it took each trained animal to find the food pellet in StQ (n = 38). Animals identified with an “x” were removed from the experiment, as their effective reinforcement time was less than half of the mean. (B) Duration of appetitive reinforcement of trained animals. (C) Plot of the animals’ center position maximum occupancy. StQ is identified with a circle in the center of the quadrant.

### Neohelice granulata shows long-term memory retention in a place preference learning task

Animals from both TR and CT groups were tested 24 hours after training session, by exposure to their cleaned corresponding arenas where all remnants of food had been removed and their behavior was evaluated for 5 minutes (Figure 5A). Contrary to their exploratory activity during the habituation, animals which had received the reinforcement in StQ for 8 minutes (TR group) spent more time in this quadrant when compared to non-reinforced crabs (CT group; p < 0.01; Figure 5B). This difference in behavior towards StQ denotes the formation of a long-term memory which lasts, at least, 24 hours after acquisition and is the result of only one training trial. In order to further characterize this type of memory, we evaluated if its strength depends on the amount of time the animal is presented with the contingency between stimuli. For this, we performed another experiment in which the total amount of time available for the animals to eat the food pellet in StQ was reduced to 2 min. When animals were able to eat for only 2 minutes, their change in behavior between Hab and TS sessions significantly differed from the behavior presented by animals which had been reinforced for an average of 8 minutes (Figure 4C). This result shows a clear dependence of the strength of the memory formed to the length of the reinforcement period.

**Figure 5.**
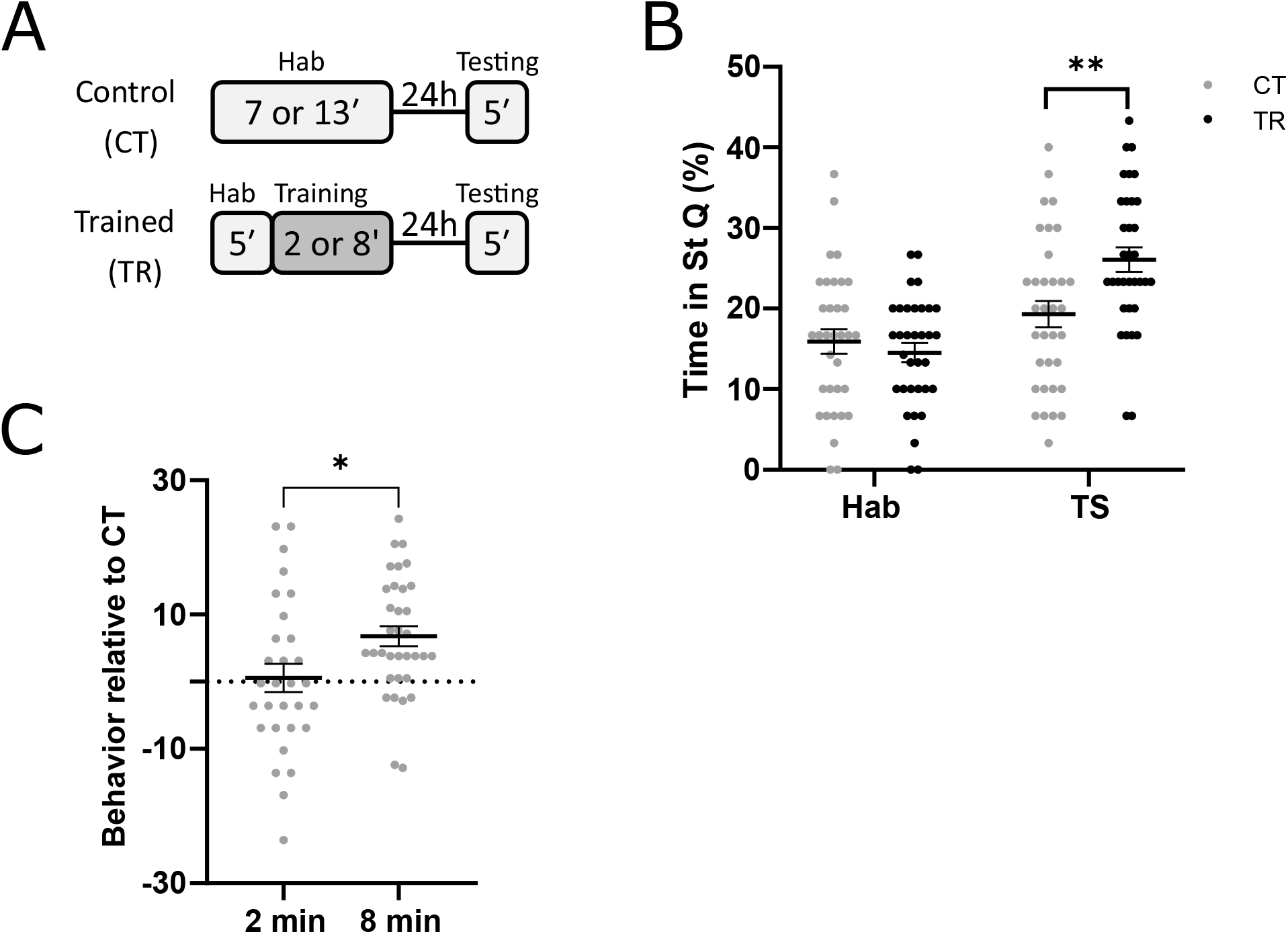
Neohelice granulata shows long-term memory retention in a place preference learning task. (A) Experimental design. (B) Percentage of time spent by crabs in the StQ, CT and TR groups n = 34 per group. Mean ± SEM. Two-way repeated measures ANOVA: session effect: F1,66 = 25.69, p < 0.001; experimental group: F1,66 = 3.41, p = 0.069; interaction: F1,66 = 7.612, p < 0.01; Contrasts: CT vs TR at Hab: p = 0.510; CT vs TR at TS: p < 0.01 (C) Group difference (TRn min-CTn min) for 2 and 8 minutes of reinforcement. Paired t-test: t62 = 2.440; p < 0.05, n (2 min) = 30, n (8 min) = 34.

### Long term retention depends on protein synthesis

Probably, the most characteristic feature of long-term memory is its dependence on de novo protein synthesis. In order to evaluate if the memory trace resulting from one training trial depends on this process, we repeated the experiment described in Figure 4A and systemically administered either cycloheximide (TRchx) or vehicle solution (TRsal) immediately after the training session (Figure 6A). Cycloheximide is a broadly used protein-synthesis inhibitor which has proven effective in several animal models, including the crab Neohelice (Fustiñana et al., 2013; Kaczer and Maldonado, 2009; Pedreira et al., 1995; Tomsic et al., 1998). Additionally, one control group was injected with vehicle solution at this time point (CTsal). When evaluating the amount of time spent by the crabs in StQ, TRsal animals showed a higher percentage relative to the total duration of TS session when compared to CTsal, which indicates the formation of a long-term memory as expected from the first experiment. Nevertheless, when animals from TRchx group were re-exposed to the arena 24h after training, they spent significantly less time than crabs which were also trained but administered with vehicle solution (TRsal; Figure 6B). Considering that a change in exploratory behavior due to protein synthesis inhibition could be actually caused by differential locomotion activities, we evaluated each animal’s total distance travelled during the TS session (Figure 6C). No significant differences were presented between experimental groups, indicating that there is no effect of cycloheximide on locomotion and that memory retention is susceptible to disruption when a protein synthesis inhibitor is injected immediately after training.

**Figure 6.**
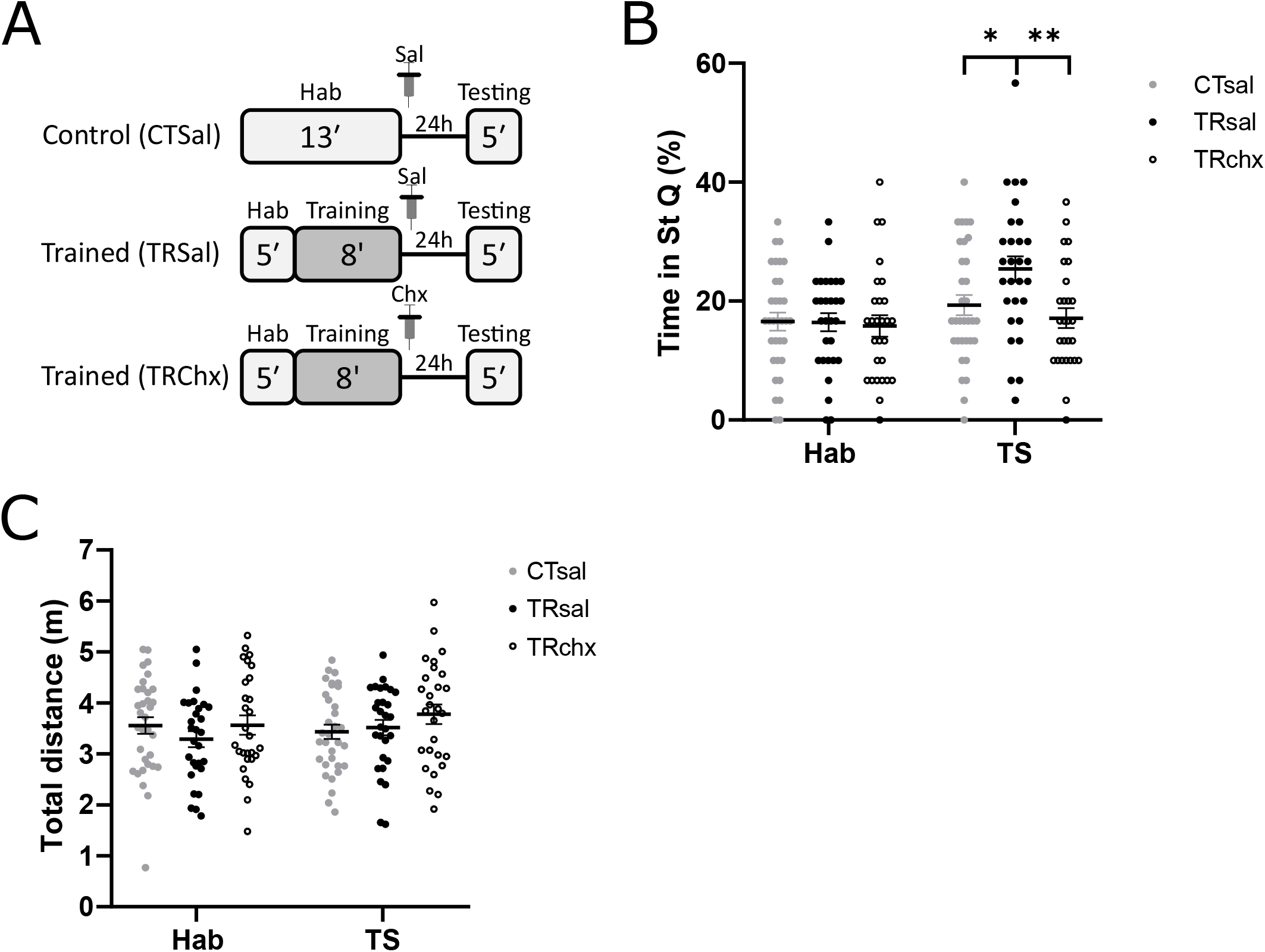
Long term memory retention is cycloheximide sensitive. (A) Experimental design. (B) Percentage of time spent by each animal in StQ. Two-way repeated measures ANOVA: session: F1,87 = 13.26, p < 0.001; experimental group: F 2,87 = 2.50, p = 0.088; interaction: F2,87 = 3.78 p < 0.05. Contrasts: within Hab: CTsal vs TRsal p = 0.99, CTsal vs TRsal p =0.99; within TS: CTsal vs TRsal p < 0.05, TRsal vs TRchx p < 0.01. CTsal n = 33, TRsal n=28 and TRchx n= 28. (C) Total distance travelled by each animal on TS session. Two-way repeated measures ANOVA: session: F1,87 = 0.90, p = 0.344; experimental group: F2,87 = 0.97, p = 0.384; interaction: F2,87 = 1.13, p = 0.329.

### Memory retention on a CS with negative valence

As seen in Figure 2B, animals could be separated by their initial predisposition to spend a greater amount of time in StQ or in the plain white portion of the arena (initial preference). During habituation, 22.25% of the crabs actually spend more time in the StQ, therefore, in order to better understand and specifically evaluate the effect of a single training trial in animals with a negative initial preference, we analyzed the behavior of the subset of crabs characterized by spending less than 25% of the habituation time in StQ (i.e., less than chance; CS-animals). We performed this type of analysis throughout 7 independent experiments, 3 of which comprised non-injected animals and 4 included animals which were injected with vehicle solution immediately after training, checking that this condition had no significant effect on the time spent in StQ (p = 0.551). As is evident in Figure 7, animals were able to overcome the initial strong aversion to the StQ and form and retain a place preference longterm memory related to a CS which was initially avoided (p < 0.01).

**Figure 7.**
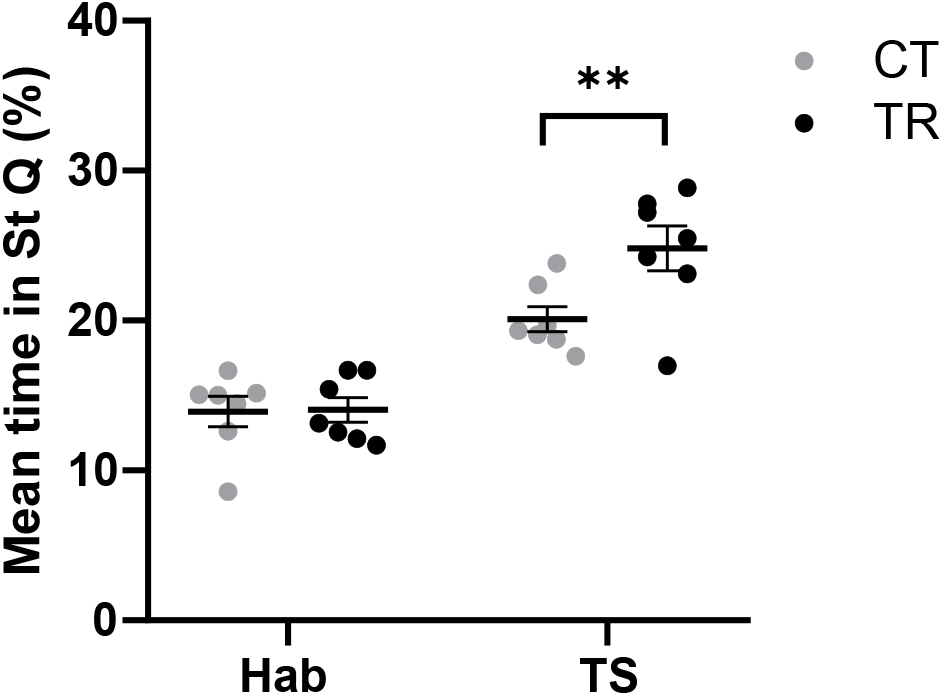
Memory retention on a CS with negative valence. Three-way repeated measures ANOVA: Session: F1,10 = 128.5, p < 0.001; Experimental group: F1,10 = 2,45, p = 0.148; Injection: F1,10 = 0.38 p = 0.551; Interaction session x injection: F1,10 = 11.39, p < 0.01. Interaction session x injection: F1,10 = 9.13, p < 0.05; Interaction injection x group: F1,10 = 0.92, p =0.359; Contrasts: CT vs TR at Hab: p = 0.947; CT vs TR at TS: p < 0.01.

## Discussion

When exposed to a novel experimental arena which comprises a plain white environment with the exception of a striped quadrant, crabs express an inhomogeneous spatial distribution in their exploratory behavior. The animals seem to react to both the quadrant delimitation in the floor and to the striped pattern in the walls of the arena during the first exposure session, presenting a thigmotactic behavior that is reduced in the target quadrant (Figure 2B). Moreover, animals presented a strong avoidance towards the target quadrant (Figure 2C and 2D). From this type of behavior, we can speculate that the striped pattern evokes some specific characteristic from the animal’s original environment (for example, the interface between the intertidal flat and the striped pattern of plant bushes), but we cannot exclude the possibility that such pattern merely consists of a negative attention attractor. Given the original avoidance fades as the crabs’ fasting period increases, an effect related to the attentional bias towards food-related stimuli is suggested, as has been previously observed in reports relating fasting periods with cognition (Benau et al., 2014). This type of modulation on avoidance behavior by the animal’s internal state resembles the behavioral changes reported in the Drosophila desensitization by hunger to bitter taste and “negative smells” (Lin et al., 2019), as well as the preponderant use of idiothetic over sensory clues for foraging and feeding in hungry flies (Kim and Dickinson, 2017). Another possible interpretation of the crab’s hunger modulation on their behavior is a shift towards the acceptance of a greater risk when exploring a novel environment, as seen by Godin and Crossman in their study on risk assessment by fish (Godin and Crossman, 1994).

The place preference memory paradigm described in the present work has the advantage of auto-administration of the reward by the animal, representing an unforced reinforcement, which could also be regulated in strength. A single training trial, which comprises a habituation period followed by the exposure to the reinforcement, results in a significant change in the avoidance behavior from the animals towards the target quadrant, which lasts, at least, 24 hours, implying the formation of a long-term place preference memory (Figure 5B). This association between the reward and the striped quadrant depends on the strength of the contingency between both stimuli, expressed by the duration in time of the reinforcement’s availability in the target zone. A short reinforcement period (2 min) is not enough to elicit the formation of a long-term memory, as an 8-min reinforcement presentation period can achieve (Figure 5C). Since other crustacean models require substantial iteration and stronger reinforcements like opioids and psychostimulants in order to form a long-term memory (Dimant, 2016; Huber et al., 2018) we think that our paradigm offers the great advantage of exploring a one-trial naturalistic reward.

On top of that, further characterization of the memory resulting from this novel place preference paradigm confirms its consolidation depends on de novo protein synthesis (Figure 6B), as has already been reported in several studies on this species (Fustiñana et al., 2013; Hermitte et al., 1999; Kaczer and Maldonado, 2009; Pedreira et al., 1995; Tomsic et al., 1998). Considering that animals were systemically administered with cycloheximide immediately after the training trial, long-term stabilization of the memory trace is likely to be impaired by the inhibition of protein synthesis, rather than its acquisition. And, since total distance travelled in the experimental arena during testing session does not differ between groups of animals which have been administered with cycloheximide or its vehicle solution, we can conclude that the difference in exploratory behavior observed between these groups is due to an effect of the drug on memory retention, rather than an unspecific effect on the crabs’ locomotion abilities.

Even though the mean behavior expressed by each of the crab’s cohorts evaluated in the experimental arena shows an avoidance towards the striped quadrant on the habituation day (Figure 2D), there still remains a minor percentage of animals expressing a preference for that quadrant (22.25% in Figure 2C). When only the initially avoiding-StQ crabs are considered in the analyses, it is possible to unambiguously evaluate the association between a conditioned stimulus with an initial negative valence and an appetitive reinforcement (Figure 7). The bases of memory retention of the association between a conditioned stimulus with an initially clear negative valence and an appetitive reinforcement could be further studied in this paradigm, opening the possibility of comparison to the animals which initially prefer the striped quadrant, together with the modulation of this preference by hunger.

In sum, we present this one-trial place preference paradigm as a classical associative conditioning, in which the crabs are able to associate a typical unconditioned stimulus (food pellet) with a clearly discernible part of the arena (striped quadrant), which comprises a nonneutral conditioned stimulus. This particular trait of the conditioned stimulus is opposed to what is traditionally expected in this type of learning paradigm, which emphasizes the importance of a neutral stimulus in order to better understand and evaluate the association between both (Pavlov, 1995). Nevertheless, it has been postulated that this type of approach is only representative of studies performed within specific laboratory conditions and fails to investigate the functional perspective of classical associative conditioning (Domjan, 2005). We strongly believe that the paradigm developed along the present work could provide a useful tool in order to study behavior in a naturalistic way, since introducing non-neutral conditioned stimuli allows evaluation of processes underlying animal conditioning as an adaptive trait.

## Author Contributions

MK and CM: Conceptualization, Methodology, Investigation, Formal analysis, Writing.

RF: Conceptualization, Resources, Methodology, Formal analysis, Writing, Project administration and Supervision.

## Acknowledgments

We would like to thank Ángeles Salles and Arturo Romano for their helpful comments and language revision which improved this manuscript, and Ángel Vidal for invaluable technical support.

